# Uncovering gene-family founder events during major evolutionary transitions in animals, plants and fungi using GenEra

**DOI:** 10.1101/2022.07.07.498977

**Authors:** Josué Barrera-Redondo, Jaruwatana Sodai Lotharukpong, Hajk-Georg Drost, Susana M. Coelho

**Affiliations:** Department of Algal Development and Evolution, Max Planck Institute for Biology, Max-Planck-Ring 5, 72076, Tübingen, Germany; Computational Biology Group, Department of Molecular Biology, Max Planck Institute for Biology, Max-Planck-Ring 5, 72076, Tübingen, Germany

## Abstract

The emergence of new genes is an important driver of evolutionary novelty. Yet, we lack a conceptual and computational approach that accurately traces gene-family founder events and effectively associates them with trait innovation and major radiation events. Here, we present GenEra, a DIAMOND-fuelled gene-family founder inference framework that addresses previously raised limitations and biases of founder gene detection in genomic phylostratigraphy by accounting for homology detection failure (HDF). We demonstrate how GenEra can accelerate gene-family founder computations from several months to a few days for any query genome of interest. We analyzed 30 genomes to explore the emergence of new gene families during the major evolutionary transitions in plants, animals and fungi. The detection of highly conserved protein domains in these gene families indicates that neofunctionalization of preexisting protein domains is a richer source of gene-family founder events compared with *de novo* gene birth. We report vastly different patterns of gene-family founder events in animal and fungi before and after accounting for HDF. Only plants exhibit a consistent pattern of founder gene emergence after accounting for HDF, suggesting they are more likely to evolve novelty through the emergence of new genes compared to opisthokonts. Finally, we show that gene-family founder bursts are associated with the transition to multicellularity in streptophytes, the terrestrialization of land plants and the origin of angiosperms, as well as with the evolution of bilateral symmetry in animals.

## Introduction

Most protein-coding genes of extant organisms descend from a small set of founder genes that were already present in the last universal common ancestor of all living systems (LUCA) (1,2). Evolutionary novelty at the molecular scale is therefore thought to be largely driven by the duplication and neofunctionalization of preexisting genetic information (3). Nonetheless, genomic studies carried out throughout the last three decades show a pervasive number of cases of genes with limited or untraceable gene homology (4–6). These taxonomically-restricted genes (TRGs) are protein-coding genes that are present in a particular evolutionary lineage with no detectable homologs in other organisms. The presence of TRGs is usually attributed to gene-family founder events, that is, the emergence of the earliest common ancestor of an extant family of protein-coding genes (7). TRGs are associated with the emergence of novel morphologies (8,9), immune defense mechanisms (10), and ecological specialization (11) across the tree of life. Proposed mechanisms that explain the birth of new gene families include neofunctionalization processes that modify the founder-gene beyond recognition (4), the differential combination and fusion of protein folds and domains that predate LUCA (12), or *de novo* gene birth from noncoding DNA (6). However, the extent to which TRGs can be attributed to gene-family founder events has been extensively debated, since the lack of traceability of a gene can also explain why some TRGs cannot be detected outside the evolutionary lineage under study (13–15). With the advent of the Earth BioGenome Project, the scientific community is reaching a stage where representative genomes will be available for a major portion of eukaryotic lineages (16). While presented as an unparalleled opportunity to study the evolutionary processes that shape genes across diverse evolutionary lineages (17), we lack a robust methodology and software solution to leverage comparative genomics at tree-of-life scale and achieve high-confidence predictions of TRG and robust assessment of gene birth events.

Genomic phylostratigraphy was initially introduced as a method to annotate gene founder events along the tree of life, often represented by a taxonomic classification (7). Inferring the relative ages of genes in a genome helps to address evolutionary questions such as the possible relation between the emergence of TRGs and lineage-specific evolutionary novelties during major radiation events (18), how ontogenetic transcriptional patterns evolve (19,20), whether new genes evolve faster compared to old genes (21), or at what rate the emergence of completely novel proteins is driven by *de novo* gene birth events (6). While conceptually powerful, several studies have questioned the detection sensitivity of the phylostratigraphic approach (13–15,22). Gene ages may appear younger than they actually are due to gene prediction errors in the target database (23). Previous approaches have also overlooked contamination or horizontal gene transfer across lineages, which can overestimate a gene’s age in a given organism (24). Furthermore, they did not estimate gene ages in terms of gene families, but assumed that dating individual genes extrapolates to the entire gene family (9,25). As such, the overall number of gene founder events is prone to be conflated by the subsequent duplication of a founder gene. Moreover, the computational burden of genomic phylostratigraphy limits its scalability. The pairwise sequence aligner BLASTP (26) (a gold standard tool to search gene homologs against sequence databases) is typically used for phylostratigraphic analyses (27,28). For phylostratigraphic applications, BLASTP has been shown to perform equally well in reporting distant homologs when compared to slower but more sensitive profile-based methods, such as HMMer (29) or PSI-BLAST (30) during the inference of gene ages (28). While faster and equally reliable as several other tools (28), a BLASTP search of a full set of organismal genes (approx. 5,000 to 40,000 genes) against currently available public sequence databases can take up to several weeks or even months, limiting its scalability to hundreds or thousands of species (17). Importantly, the greatest caveat of genomic phylostratigraphy is that small and fast-evolving genes are often mis-annotated as young genes due to homology detection failure (HDF), *i*.*e*, the inability of pairwise local aligners to trace back distantly-related homologs only due to neutral sequence divergence which results in spurious patterns of TRG birth (13,15). Overall, these caveats undermine the power of the original phylostratigraphic method, motivating several authors to propose key methodological improvements to obtain a more reliable estimate of gene-family founder events (9,15,23,24,28,31,32).

Here, we present a conceptually redesigned founder-gene inference method that employs the superior computational speed of the protein aligner DIAMOND (17). This method draws from the principles of genomic phylostratigraphy (7) to accurately pinpoint the evolutionary origin of gene families, but extends its initial scope to account for founder gene-family origination events and sufficient homology detection sensitivity. We apply this method to the reference genome of *Saccharomyces cerevisiae* (33) to benchmark its scalability and accuracy when searching against thousands of target genomes. We use our methodology to revisit the putative pattern of TRG emergence associated with important evolutionary events in plants and animals, such as the transition to multicellularity in animals or the terrestrialization event of plants (4). We also explore whether analogous TRG patterns are also present in Fungi. Furthermore, we calculate and investigate the gene age maps of 30 genomes across vastly different lineages within three different eukaryotic kingdoms to test whether accounting for HDF changes the observed patterns of TRG emergence (15), as some radiation events may lead to a strong signal of gene untraceability (34). Finally, we investigate the presence of ancient protein domains within these TRGs to evaluate the relative contribution of gene duplication and domain reshuffling in TRG emergence compared to *de novo* gene birth.

## Materials and Methods

### Addressing the previous shortcoming of genomic phylostratigraphy with GenEra

GenEra address all major limitations and scalability of previous phylostratigraphic approaches while expanding its functionality by implementing a DIAMOND-fuelled method to detect gene-family founder events (Fig. 1A). The pipeline can be used on the full set of genes from any species whose taxonomy is included in the NCBI database (35). We provide this pipeline as an open source command line tool called GenEra (https://github.com/josuebarrera/GenEra).

**Figure 1.**
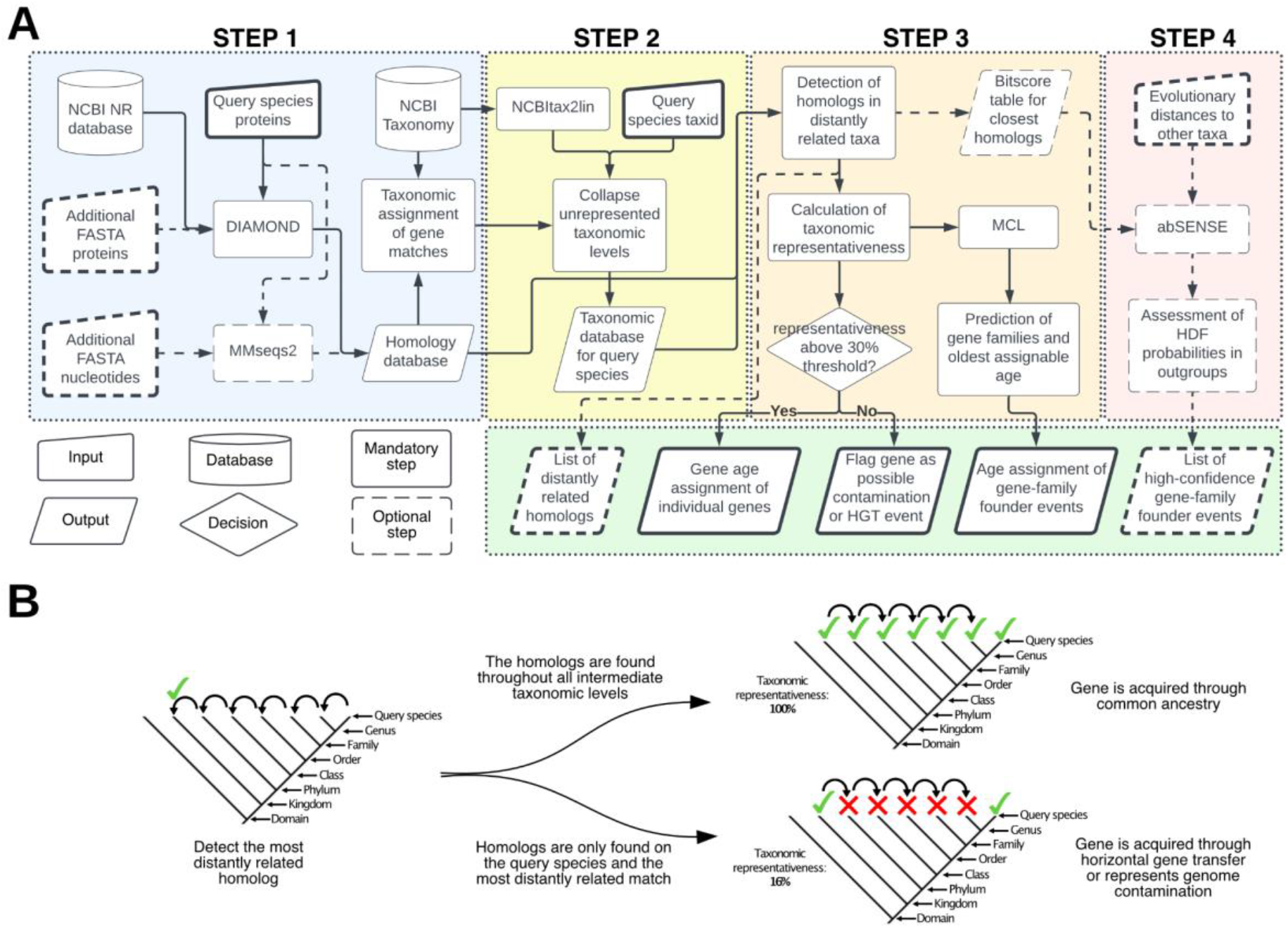
Gene-family founder detection framework implemented in GenEra. Overview of the pipeline for sensitive founder-gene family detection across the tree of life. A) Flowchart of the command-line tool GenEra. Solid arrows/elements represent the mandatory steps in the pipeline, while the dashed arrows/elements represent optional steps to enrich the results. B) Graphic representation of the rationale behind the taxonomic representativeness score (see Methods). GenEra first performs a taxonomic trace-back to determine the most distantly related homolog to a query species, and then tracks back the presence of homologs in all the intermediate taxonomic levels, which helps to detect putative contaminants in the query proteome, horizontal gene transfer events between increasingly distantly related taxa, or false positive matches.

The first step of GenEra replaces BLASTP with the ultra-fast and sensitive protein aligner DIAMOND, which has recently been extended for ultra-sensitive gene similarity assessments at tree of life scale (17). By default, BLAST and other sequence search algorithms limit the maximum number of top sequence hits that are reported in the analysis to the 500 best hits, which is an often overlooked limitation that hinders the extent by which genes can be traced back to distantly related taxa. With exponentially growing sequence databases covering hundreds of thousands of species, 500 top hits can at best cover only 500 different subject species, thereby losing a significant proportion of age-assignable information. Using DIAMOND instead of BLAST allows us to build a customized list of pairwise alignments against the entire NCBI non-redundant (NR) protein database, which harbors tens of thousands of genomes, alongside other user-defined protein datasets with an unlimited amount of sequence hits, generating results up to 8,000 times faster than BLASTP-based approaches while reaching the same level of accuracy (Fig. S1) (17). We established an e-value threshold below 1e^-5^ for a sequence hit to be considered a reliable true positive. The choice of this threshold was based on an extensive threshold-robustness study to test the influence of a diverse range of e-values on gene age assignments with the ultimate aim to determine the most robust e-value threshold when running GenEra in default mode. Indeed, a less stringent threshold does not improve the age assignment of genes and may lead to an increased rate of false positive age assignments, given the size of the NR database, while more stringent thresholds lead to an overestimation of TRGs (Fig. S2). Another issue raised for genomic phylostratigraphy is that spurious genome annotations or comparison between annotations with different levels of quality and accuracy can overestimate the proportion of recently-evolved proteins in the analysis (23,31). To address this short-coming, GenEra is able to include an additional protein-against-nucleotide search through MMseqs2 (36) with its most sensitive parameters (s = 7.5) to reconfirm gene age assignments with an annotation-free approach solely based on six-frame alignments. Using GenEra with genome assemblies in parallel with protein annotations (Table S1) significantly reduces the number of TRGs in the youngest taxonomic levels, from the species-level up to the genus-level age assignments, but older taxonomic levels seem largely unaffected when including protein-against-nucleotide data (Fig. S3).

The second step of GenEra employs NCBItax2lin (available via https://github.com/zyxue/ncbitax2lin) to generate a lineage database that is used to associate the NCBI Taxonomy ID in the list of DIAMOND pairwise alignments with their hierarchical taxonomic identity in the NCBI Taxonomy database. The NCBI Taxonomy is a curated database that reflects the current knowledge of the relationships between all known organisms (35). Hence, each taxonomic level in the lineage database corresponds to a monophyletic group in a species tree. This allows GenEra to determine the evolutionary relationship between the matching genes from the sequence database and the query species. The lineage database that is generated by NCBItax2lin is not arranged in a hierarchical order, given that the taxonomic ranks are usually asymmetrical between different lineages in the NCBI Taxonomy database (37). Thus, GenEra retrieves the correct taxonomic order from the NCBI server to rearrange the lineage database in a hierarchical order, in accordance with the taxonomic levels that are reported in the NCBI for the query species. Given the historical scopes and interests of the scientific community during the era of high-throughput sequencing, current genomic databases are still biased toward having sequencing data for certain groups of organisms, while having no publicly available genomes for others (38). This complicates the detection of gene-family founder events, since having genomic data is required to assign a gene to a certain age in a reliable and systematic manner. To address this issue, GenEra searches the entirety of sequence matches that were retrieved with DIAMOND and only retains the taxonomic levels for which at least one representative species matches more than 10% of the proteins in the query species for further analyses. This threshold was empirically established to exclude the organisms in the NR that are represented by only a few genes and not by genomic data (Fig. S4).

The third step of GenEra performs a taxonomic trace-back to determine the most distantly-related lineage that matches each gene of the query species (Fig. 1B). Once the most distant homolog for a query protein is found, the pipeline calculates a taxonomic representativeness score to estimate the reliability of assigning a gene age based on this sequence match. The rationale for this procedure is to address another limitation of the original genomic phylostratigraphy, where the most distant hit was not reconfirmed at higher taxonomic levels but rather assumed, which created a systematic bias when dealing with contamination and horizontal gene transfer events. GenEra reconfirms hits at higher levels using a taxonomic representativeness metric (*L*) which is calculated as the presence of homologs in at least one representative species for each of the intermediate taxonomic levels between the most distantly-related lineage and the query species (Fig. 1B). The number of internode taxonomic levels with representative gene homologs (*RP*) is divided by the total number of taxonomic steps that separate the most distantly-related match from the gene of the query species (*AP*), while excluding the youngest taxonomic level (usually the species level), since the presence of the gene in the query species already confirms its representativeness at that level:

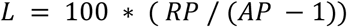

This gives a taxonomic representativeness score *L* with a scale from 100 to 100*(1/(*AP* - 1)), which helps to flag genes that are only present in the query species and in other distantly related taxa (Fig. 1B). Genes with a low taxonomic representativeness are discordant with the concept of synapomorphy (39), where a homologous character (in this case, a gene) should be inherited to all the taxa that share a common ancestor. However, secondary losses of inherited genes should be expected to happen throughout the tree of life. Thus, the taxonomic representativeness score can be influenced by gene loss events in the genomes that act as representatives in the intermediate taxonomic levels, or due to the availability of only scarce and low-quality genomic data at certain taxonomic levels. To address this issue, we established a relaxed taxonomic representativeness threshold of 30%, so that only genes with a particularly low score are flagged as putative horizontal gene transfer events, contaminant sequences in the assembly that do not belong to the query species, or false positive matches against the database (Fig. S5). This score is reported for every gene in the query species, and the user can also establish a custom threshold that is appropriate for the dataset and taxon of interest.

GenEra can optionally report the best sequence hit (as defined by its bitscore) that can be assigned to the oldest taxonomic level for each query gene. This feature helps users to identify erroneous gene age assignments due to false positive matches, and to manually evaluate genes with a low taxonomic representativeness. This feature also helps to identify candidate non-coding sequences from which potential *de novo* TRGs could have emerged when implementing a protein-against-nucleotide search.

Once all the genes in a query species have been assigned to a certain age, GenEra performs an all-vs-all DIAMOND search of the query proteins against themselves to detect paralogs within the genome of the query species. The e-values of the all-vs-all DIAMOND search are normalized through a negative log10 transformation, and are subsequently used for a clustering analysis with an inflation value of 1.5 to predict gene families using MCL (40). GenEra uses the oldest assignable gene age for each of these gene clusters to estimate the number of gene-family founder events throughout the evolutionary history of the query species.

GenEra has a fourth additional step to assess whether the gene age assignment of the query genes can be explained by HDF. Bitscores obtained through pairwise sequence alignments have been shown to decay exponentially as a function of evolutionary distance (15). Given enough data points, one can calculate the expected bitscore for a given gene in a distantly related species when such gene is not detected, and hereby calculate the probability of not finding this gene as a consequence of bitscore decay alone (15). When GenEra is given a list of pairwise evolutionary distances (*e*.*g*., substitutions per site in a phylogenetic tree) between the query species and other taxa in the database, it searches for the closest homolog in these species, which are defined as the highest bitscore matches to each of the query genes. GenEra uses the bitscore of these genes to calculate HDF probabilities using abSENSE (15) for all the species that lack any traceable homolog to each query gene in the target species. GenEra can use these probabilities to test the null hypothesis of untraceable homology for each gene that is assigned to a given taxonomic level. The ability of GenEra to test HDF for each taxonomic level is dependent on the taxonomic sampling that is given by the user, which is determined by the taxonomic sampling of the phylogeny that is used to calculate the evolutionary distances. Hence, the use of phylogenies at different taxonomic levels can be used by GenEra to test for HDF in gene-family founder events at different evolutionary scales. Once a gene is assigned to a certain age, GenEra analyzes the HDF probability of the closest species (as defined by their evolutionary distance to the query species) that belongs to the next taxonomic level, and labels the gene age assignment as a high-confidence gene founder event whenever the HDF probabilities fall below 0.05 in the outgroup (Fig. S6). Thus, GenEra can make an informed decision on whether the gene age assignments can be explained by gene-family founder events or through sequence divergence alone that make these genes untraceable given their size and substitution rate (27).

### Assessment of gene-family founder events in 30 eukaryotic genomes

We downloaded representative genomes of plants, animals and fungi from the Uniprot reference proteomes to study the patterns of gene-family founder events throughout these major eukaryotic lineages by using GenEra (Table S2). We chose 10 representative taxa across the taxonomic diversity of each of these three kingdoms to revisit the previously observed peaks in gene-founder events associated with the diversification of animals and land plants (4), and to determine whether this same pattern arises in Fungi. We collapsed the Eumetazoan taxonomic level (i.e. all animals excluding Porifera) from our animal analysis, since recent evidence indicates that Eumetazoa is paraphyletic (41).

We extracted the evolutionary distances from previously reported phylogenies using the ape package in R (42) to calculate HDF probabilities at different taxonomic levels and evaluate the proportion of gene families that can be confidently assigned to gene-family founder events. For the Fungi kingdom, we used 81 evolutionary distances from a maximum likelihood tree (43) encompassing several evolutionary distances from our 10 target genomes (Table S3), including *Fonticula alba* and other opisthokonts as outgroups to test gene-family founder events up until the Fungi level. For Metazoa, we used 43 evolutionary distances from a posterior consensus bayesian tree (41) comprising a large portion of the animal phyla (Table S4) and which includes *Monosiga brevicollis* and *Salpingoeca rosetta* as outgroups (44) to test gene founder events at different taxonomic levels up to Metazoa. For Embryophyta, we used 61 evolutionary distances (Table S5) from a posterior consensus bayesian tree (45) that incorporates several plant genomes, as well as green algae and red algae, which helped us test gene-family founder events up until the Viridiplantae level. All the gene families who had HDF probabilities < 0.05 in the closest outgroup were considered as high-confidence TRGs that resulted from gene-family founder events.

## Results

By improving genomic phylostratigraphy with a gene family clustering strategy and HDF tests, we were able to estimate the number of putative gene-family founder events throughout the plant, animal and fungal lineages. We used 10 genomes for each of these lineages to evaluate the common patterns of putative gene founder events that have been previously described using genomic phylostratigraphy with single genomes (4). All our results are available as supplemental data.

Before calculating HDF probabilities, we found a consistent overrepresentation of putative gene-family founder events at the taxonomic levels that correspond to the crown node of land plants, animals and fungi (Fig. 2). These gene age peaks were observed independently of the taxonomic lineage from which the species belong within their kingdom, revealing a common evolutionary signal. We found no evidence of whether this convergent pattern was correlated with the number of available genomes in the database at those taxonomic levels, as these levels can have a vastly different number of representative genomes depending on the species that is analyzed (Table S6).

**Figure 2.**
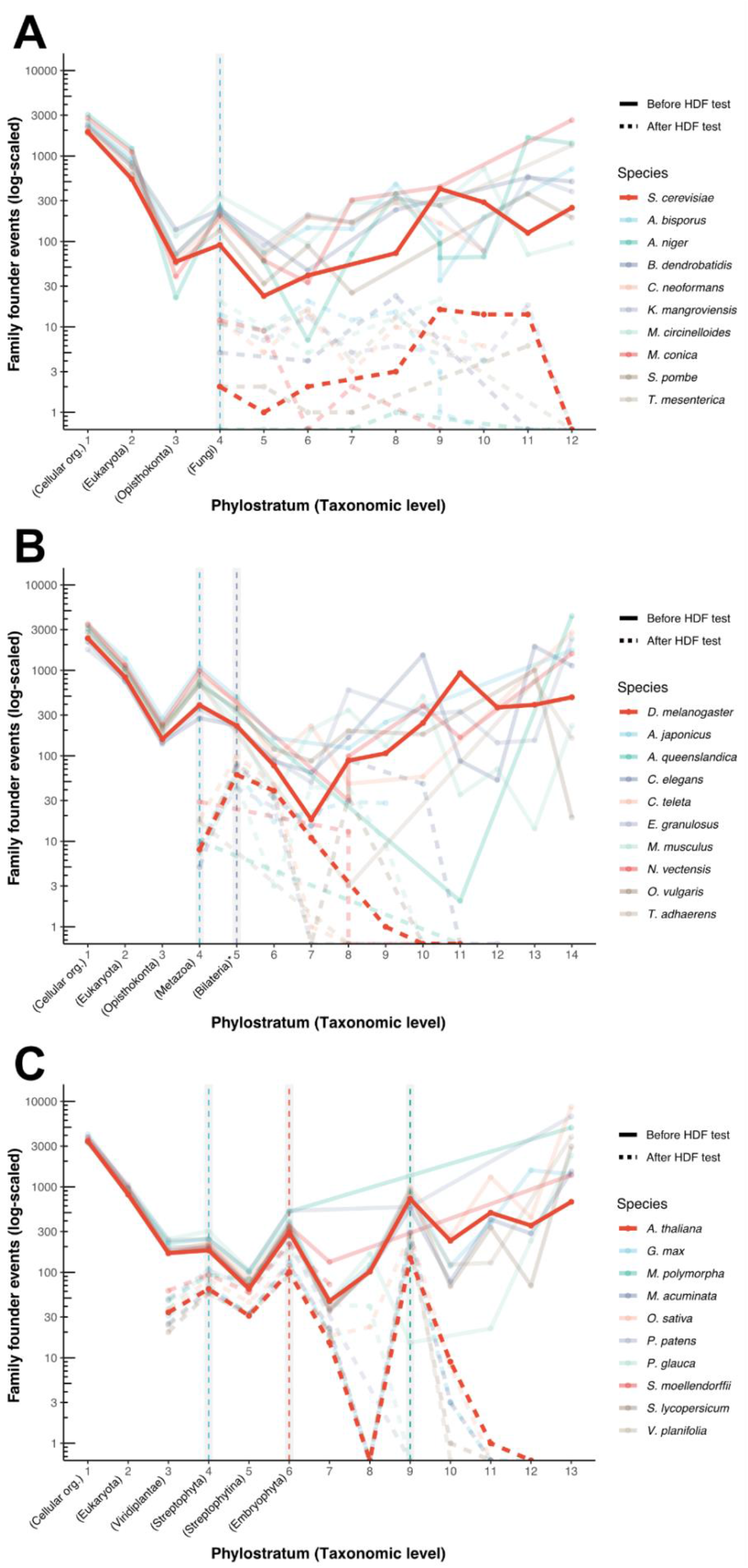
Detection of gene-family founder events at major evolutionary transitions in plants, animals and fungi. Overlapping plots of gene-family founder events before and after accounting for HDF (solid lines and dashed lines, respectively). The taxonomic hierarchies that are shared between all the species are named in the X axis, while the taxonomic levels that differ between species are just ranked by number. A) Fungi genomes with *S. cerevisiae* as the representative species in the plot. The taxonomic level leading to the emergence of Fungi exhibits a burst of gene-family founder events before the HDF test (dashed blue line), but all the common patterns are lost after accounting for HDF. B) Animal genomes with *D. melanogaster* as the representative species in the plot. The taxonomic level leading to the emergence of Metazoa also shows a burst of gene-family founder events before the HDF test (dashed blue line). The Metazoa burst fades after accounting for HDF, but the taxonomic level of Bilateria exhibit a burst after the HDF test for all bilaterian animals (\**i*.*e*., excluding *N. vectensis, T. adhaerens* and *A. queenslandica*; dashed grey line). C) Plant genomes with *A. thaliana* as the representative species in the plot. Plants genomes display a consistent pattern of gene-family founder events before and after accounting for HDF, with gene-family founder bursts associated with the emergence of multicellularity (Streptophyta, blue dashed line), the conquest of land by plants (Embryophyta, red dashed line) and the origin of flowering plants (Magnoliopsida, green dashed line).

However, the patterns of gene-family founder events considerably change after filtering the dataset by HDF probabilities. The total number of putative founder events diminished between one and two orders of magnitude in all the analyzed species after retaining only high-confidence gene ages that could not be explained by HDF. Fungi lost any discernible pattern of gene emergence that could be traced back to a particular evolutionary transition after accounting for HDF, including the putative TRG overrepresentation at the kingdom level (Fig. 2A). Likewise, the signal associated with the emergence of Metazoa is lost in the high-confidence gene-family founders, but the transition to bilateral symmetry (Bilateria) is consistently enriched in high-confidence gene-family founder events on all the bilateral animals in our dataset (Fig. 2B). We analyzed the biological function of these TRGs by looking at the gene annotation of *D. melanogaster*. We detected the emergence of the Ninjurin A-C genes, the *disconnected* gene, the Dampened gene and the gene family composed of the Gurken, Keren and spitz genes.

The patterns of gene-family founder events in plants remained fairly consistent despite predicting a vastly smaller amount of founder events. The most consistent bursts of gene-family founder events in plants were found in Streptophytes when green algae transitioned to complex multicellularity (46), in embryophytes when plants conquered the land (47), and in angiosperms, when plants evolved flowers (48) (Fig. 2C). We inspected the gene annotations of *A. thaliana* to evaluate the biological function of these TRGs.

Some of the successful gene-family founder events that were identified as high-confidence Streptophyta TRGs include a family of Basic Helix-Loop-Helix (bHLH) transcription factors (49), the COBRA-like gene family that act as key regulators of cell-wall expansion in the meristems (50), a family of auxin canalization proteins that regulate plant growth through auxin transport (51) and the BRASSINAZOLE-RESISTANT family of transcription factors that modulate brassinosteroid signaling in plants (52).Surprisingle, ARABIDILLO and the ULTRAPETALA gene families appear to be Streptophyta TRGs, with putative homologs in the charophyte algae *Klebsormidium nitens* (GAQ84482.1 and GAQ90507.1, respectively).

The high-confidence gene-family founder events that were linked to the emergence of embryophytes include a family of F-box/kelch-repeat proteins that regulate the biosynthesis of phenylpropanoids (53), the group 2 of late embryogenesis abundant (LEA) proteins that are involved in plant response to osmotic and oxidative stress due to desiccation (54), two groups of bHLH transcription factors (49), a gene family that contains MORPHOGENESIS OF ROOT HAIR 6 (MRH6), a gene family that contains Piriformospora indica-insensitive protein 2 (PII-2), the SOSEKI gene family that regulates cell polarity in early plant development (55) and the LONGIFOLIA gene family, involved in leaf development (56).

Within the gene-family founder events in angiosperms, we found class III of Ovate family proteins (OFP) and the family of paclobutrazol resistance (*PRE*) genes. However, most of the TRGs in this taxonomic level belong to genes that are uncharacterized in *A. thaliana*. The founder event of the MADS-box gene family was detected in LUCA.

## Discussion

Gene founder events facilitate evolutionary innovations. Determining the timing of these events is therefore important for evolutionary research. Such inference is not trivial, since previous attempts to estimate TRG birth have overlooked the effects of HDF (4,7–9,18). While these initial efforts were useful for investigating more general processes of evolution, such as the assessment of transcriptome age during development (19,20), they lack the detection sensitivity to decouple founder events of entire gene families from patterns of gene untraceability. For this reason, we developed GenEra to provide the community with a sensitive and computationally optimized approach for gene-family founder detection across the tree of life. To demonstrate the versatility of GenEra, we analyzed 30 genomes from plants, animals and fungi to capture the broad diversity of gene-family founder events in these lineages. We show that GenEra is applicable to any eukaryotic genome and we provide extensive documentation to facilitate its swift adoption in the life science community.

The origin of TRGs has sparked important debates over the last decade regarding the processes of gene birth (4–6,13–15,22,27). A high proportion of gene age assignments in our dataset could be explained by HDF, as previously reported (13–15). It is important to acknowledge that gene age assignments that fail the HDF test should not be interpreted as not belonging to their estimated taxonomic level, but rather that we cannot reject the null hypothesis of untraceable homology in more distantly related lineages (15). This is particularly true for short and fast-evolving genes, which are prone to fail the HDF test (15), but which are also more likely to have arisen recently, given that previously validated *de novo* genes are usually shorter and have fewer exons compared to old genes (11,57). Nevertheless, the probability of proteins to independently acquire similar tertiary structures *de novo* is astronomically small, given that the number of possible amino acid configurations of a 100-residue protein that is considered “small” (58) (20^100^ configurations) is bigger than the estimated number of atoms that are contained in the observable universe (∼10^82^ atoms) (2). Hence, finding conserved motifs and domains outside the boundaries of TRGs should be regarded as compelling evidence to discard *de novo* birth scenarios. The vast majority of the high-confidence TRGs we detected contain highly conserved protein domains and motifs that are consistently found throughout the tree of life. Such is the case for the bHLH motif, which can be found in transcription factors of all eukaryotes (59), the DIX domain in SOSEKI genes that are also conserved throughout eukaryotes (55), the ARMADILLO repeat domain in ARABIDILLO genes that can be found in animals (60) or transmembrane domains found throughout all cellular organisms (61). These TRGs cannot be explained by HDF (13,15), nor through *de novo* gene birth, as previously suggested (5). Our observations support the idea of gene duplication (3) and of protein modularity, where gene-founder events result from the differential fusion of pre-existing folds and domains (12), whose tertiary structure acquired the property to fold during the postulated era of the RNA and peptide world (1,2). These domain-containing TRGs were coincidentally found as multi-copy gene families, suggesting that old protein folds and domains were already optimized by natural selection to perform their biological activity (1,2), ensuring the evolutionary success of these TRGs. Despite the minor role of *de novo* gene birth in TRG emergence, the study and validation of successful *de novo* founder events should be of particular interest for evolutionary research, as these events can help us uncover the foundations of evolutionary novelty at the molecular level (62).

Our results before the HDF test retrieved analogous peaks of gene age assignments in plants and animals that have been previously described by Tautz and Domazet-Lošo (4) and could extend their insights by detecting a kingdom-level peak in Fungi. The consistency of these peaks throughout several species with vastly different evolutionary histories and biological traits (*e*.*g*., free living organisms and parasites, unicellular and multicellular fungi, plants with haploid-dominant and diploid-dominant life cycles, bilateral-symmetric and non-bilateral-symmetric animals) and the lack of a correlation between the database and gene age assignment (Fig. S7) points towards a biological basis of such a convergent pattern. However, the biological interpretation of TRG patterns should be taken with caution. These TRG peaks have been previously interpreted as bursts of genomic novelty that have accompanied some important diversification events throughout the evolutionary history of these lineages (4,9), but we found that the overrepresentation of TRGs at the emergence of animals and fungi disappears after accounting for HDF, suggesting that these peaks may be driven by untraceable homology beyond those taxonomic levels (15), rather than gene-family founder events or any other source of molecular novelty.

The emergence of animals and fungi are associated with their independent emergence of multicellularity (63) and the diversification bursts that followed this key evolutionary innovation (34). Diversification events have long been known to correlate with molecular substitution rate accelerations (64–66), even though the exact causal relationship between both phenomena remains underexplored (67). If substitution rates are correlated with diversification events, we would expect a large proportion of the genes in the genome to become untraceable beyond these major diversification bursts. Accordingly, our analyses show a pattern of gene untraceability that is linked to the emergence and the diversification bursts of these two eukaryotic kingdoms. Therefore, we propose that these gene age assignment peaks are driven by substitution rate accelerations that were linked to the diversification bursts that accompanied these major evolutionary transitions in animals and fungi. Although gene emergence likely played an important role during these evolutionary transitions in the tree of life, our results indicate that gene-family founder events may not be as pervasive in the emergence of evolutionary novelties such as multicellularity in opisthokonts compared to the co-option of ancient gene families that already existed in LUCA, such as transcription factors, cell-adhesion proteins and cell-signaling genes, which likely drove biological novelty through novel regulatory pathways (68,69). Furthermore, recent studies suggest multiple origins of multicellularity in Fungi through vastly different evolutionary processes compared to animals or plants (69). This likely blurs any common pattern between molecular innovations and the transition to multicellularity in fungi. A more in-depth analysis of fungal genomes might elucidate key gene-family founder events in this eukaryotic lineage.

We found a consistent overrepresentation of gene-family founder events in Bilateria. The emergence of Bilateria is defined by a change in developmental patterns that resulted in the evolution of bilateral symmetry. Among our reported gene-family founder events, we found Gurken, spitz and Dampened as Bilateria TRGs. These genes are all involved in the establishment of the anterior-posterior and dorsal-ventral polarities and neurogenesis during development (70–72). Likewise, the protein *disconnected* is involved in the formation of the nervous system and the connection of the visual nerve to the brain (73).

Our results show that three major evolutionary transitions in plants are associated with the evolution of entire new gene families. The observed pattern of TRG birth in plants is conserved even after accounting for HDF, suggesting that plants are prone to evolve novel traits through the emergence of new genes. The frequency of gene-family founder events in plants could be driven by the propensity of their genomes to undergo structural rearrangements and whole-genome duplications (74). Our results are consistent with an orthogonal approach by Bowles et al., where they find an independent burst of gene novelty in the branch leading to Streptophyta and Embryophyta (8), even though that study did not account for HDF, which likely inflated the amount of predicted TRGs at those taxonomic levels. Streptophytes, which include land plants and charophytes, have been proposed to share a common emergence of complex multicellularity (8,46). Complex multicellularity has been linked with the expansion of transcription factors, the emergence of an internal communication system between cells (63) and, in the case of plants, the emergence and expansion of cell-wall remodeling proteins (46). Coincidentally, our analysis detected gene-family founder events in bHLH transcription factors (49), BRASSINAZOLE-RESISTANT transcription factors (52), COBRA-like genes (50) and auxin canalization proteins (51). Furthermore, the emergence of auxin canalization proteins and BRASSINAZOLE-RESISTANT genes likely contributed to the establishment of an internal communication system between cells in multicellular streptophytes through the regulation of the basic hormone-receptor systems that predate the evolution of multicellularity (75). We found a putative ULTRAPETALA and ARABIDILLO homologs among charophyte algae, even though these gene families were previously reported as embryophyte and angiosperm TRGs, respectively (60,76). ARABIDILLO genes have been co-opted to modulate different developmental processes in plants through abscisic acid signaling (60), while ULTRAPETALA genes intectacts with the trithorax group of angiosperms to coordinate flower development through chromatin-dependent transcriptional regulation (76). If the homologs found in *Klebsormidium nitens* are reliable, this would suggest an early role of ULTRAPETALLA and ARABIDILLO homologs in streptophyte evolution (77).

The evolution of land plants (Embryophyta) is intertwined with an increased morphological complexity compared to other streptophytes. The emergence of SOSEKI genes probably conferred plants with cell-polarization mechanisms to ensure the correct development of complex multicellularity (55). The LONGIFOLIA gene likely played an additional role in the emergence of complexity in land plants through the development of leaves (56). The emergence of embryophytes has also been associated with the emergence of several defense mechanisms to cope with the abiotic stresses that characterize the transition from water to land, such as ultraviolet (UV) radiation, drought and temperature fluctuations (47). Accordingly, we found a F-box/kelch-repeat gene-family founder event in Embryophyta, whose gene members regulate phenylpropanoid biosynthesis (53). The production of phenylpropanoids has long been recognized as a crucial adaptation in plants that allowed them to survive the effects of UV radiation in land (47). The emergence of group 2 LEA proteins would have helped plants to cope with drought stress (54) as they transitioned from water to land. The role of rooting structures and its association with mycorrhizal fungi have also been proposed as important innovations in land plants

(47). We detected two bHLH groups in our analysis, which have been shown to coordinate the development of rhizoids and roots in plants (78). We also detected MRH6 as an embryophyte TRG, which is involved in root hair development (79). Furthermore, PII-2 is known to promote plant growth and seed production through its interaction with the mycorrhizal fungus *Piriformospora indica (80)*, whose detection as an embryophyte TRG supports the role of plant-fungus interactions in the transition from water to land (47).

The emergence of flowers and fruits are major evolutionary innovations in angiosperms that changed the ecological dynamics of terrestrial life (48). Many genes that regulate flower development are known to belong to old gene families, such as the MADS-box genes (81). Accordingly, our analysis retrieved the founder event of MADS-box genes in LUCA. However, our results also detected the founder event of the class III OFP genes, which are also involved in the development of fruits (82). Most of the founder events we detected angiosperms belong to uncharacterized genes with unknown biological activity. The experimental study of these TRGs should shed light on the evolution of flowering plants.

The consistency of our results with previous studies on the emergence of these widely studied evolutionary transitions highlights the power of GenEra to accurately detect molecular innovations in three different eukaryotic lineages. Our method is expected to be a useful resource to detect novel gene-family founder events on other important transitions throughout the tree of life.

## Acknowledgments

The authors gratefully thank Caroline M. Weisman for her helpful comments on how to analyze and interpret HDF probabilities. We thank the Max Planck Computing and Data Facility for access to and support of the HPC infrastructure, as well as the BMBF-funded de.NBI Cloud within the German Network for Bioinformatics Infrastructure (de.NBI) (031A532B, 031A533A, 031A533B, 031A534A, 031A535A, 031A537A, 031A537B, 031A537C, 031A537D, 031A538A).

## Author contributions

JB-R, JSL, H-GD and SMC conceived the study. JB-R designed and implemented the software. JB-R and JSL analyzed the data. JB-R, JSL, H-GD and SMC interpreted the data. JB-R, JSL, H-GD and SMC wrote the manuscript.

## Supplementary figures

**Figure S1.**
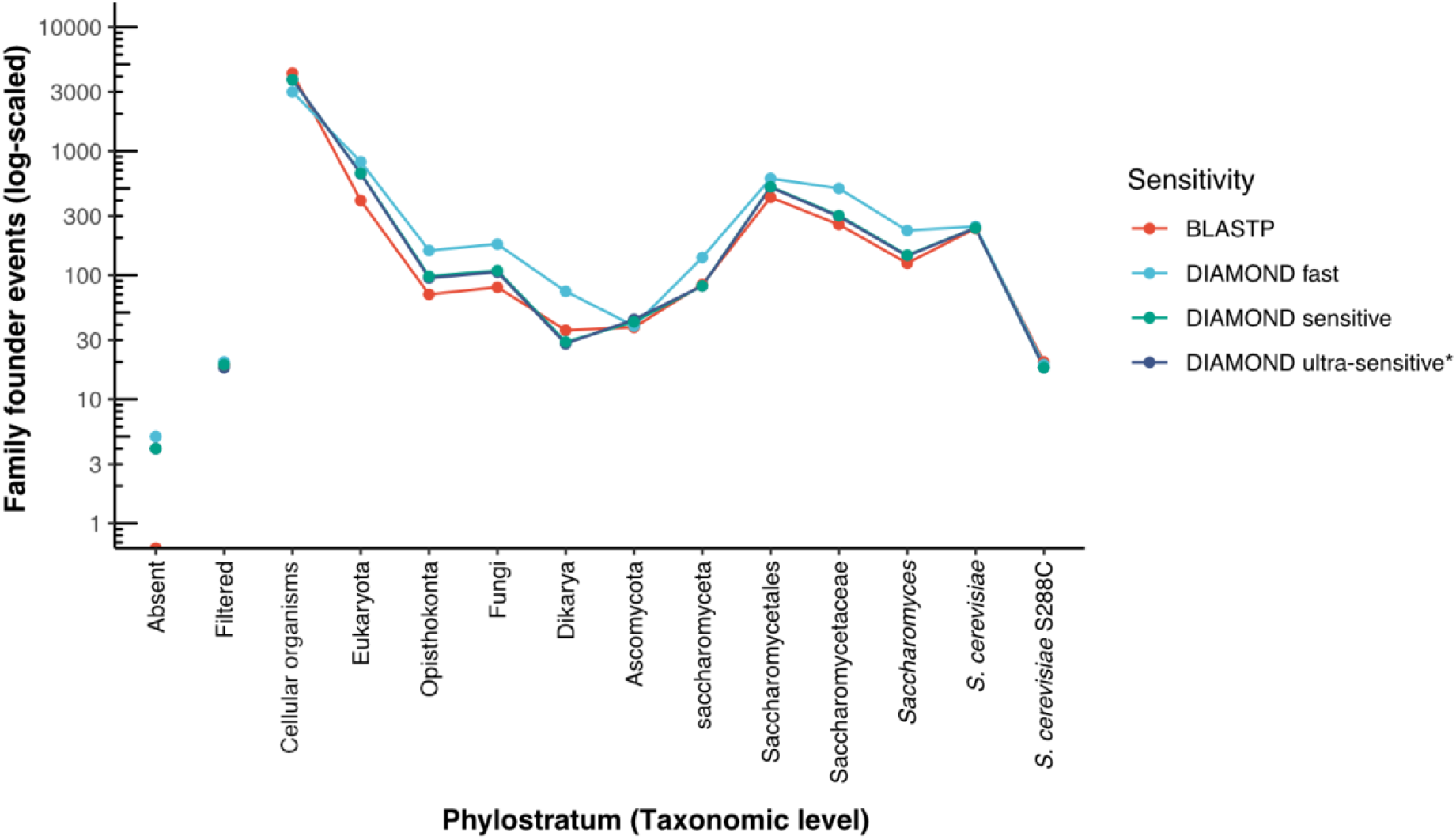
Impact of DIAMOND search sensitivity level on gene age assignment in *S. cerevisiae* while using BLASTP as reference. DIAMOND in ultra-sensitive and sensitive mode generates a similar pattern of gene age assignment as the gold-standard of BLASTP while using the same e-value threshold of 1e^-5^. DIAMOND in fast mode assigns younger ages to genes compared to more sensitive modes as a consequence of not finding as many distantly related homolog genes in the database. The search sensitivity level does not influence the number of genes that are filtered through the taxonomic representativeness threshold (filtered; values below 30% taxonomic representativeness), and has a negligible effect on the number of genes that fail to match themselves through pairwise alignment (absent). We established ultra-sensitive as the default sensitivity mode for GenEra (asterisk).

**Figure S2.**
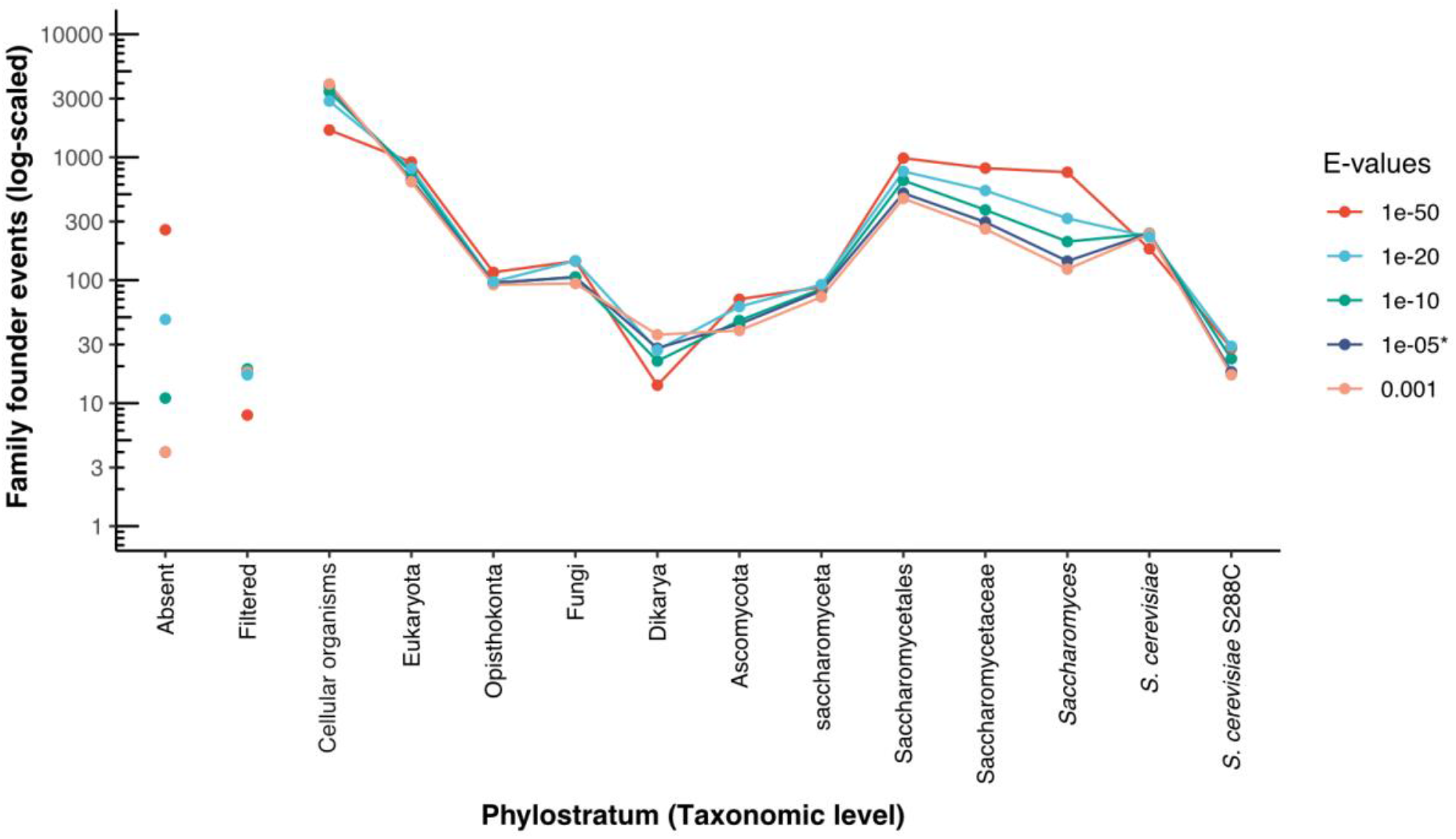
Impact of e-value thresholds on gene age assignments in *S. cerevisiae*. The patterns of gene age assignment remain largely unaffected between a premisive e-value threshold of 1e^-3^ and a more stringent threshold of 1e^-5^. Using more stringent thresholds (1e^-10^ or lower) leads to an overrepresentation of TRGs at younger taxonomic levels. Lower e-value thresholds also increase the amount of genes whose self-alignment cannot be detected (absent), likely increasing the amount of false negative matches in the database. The e-value threshold also has a small influence on the proportion of genes that are discarded through the taxonomic representativeness threshold (filtered; values below 30% taxonomic representativeness). We established a default e-value threshold of 1e^-5^ for GenEra (asterisk).

**Figure S3.**
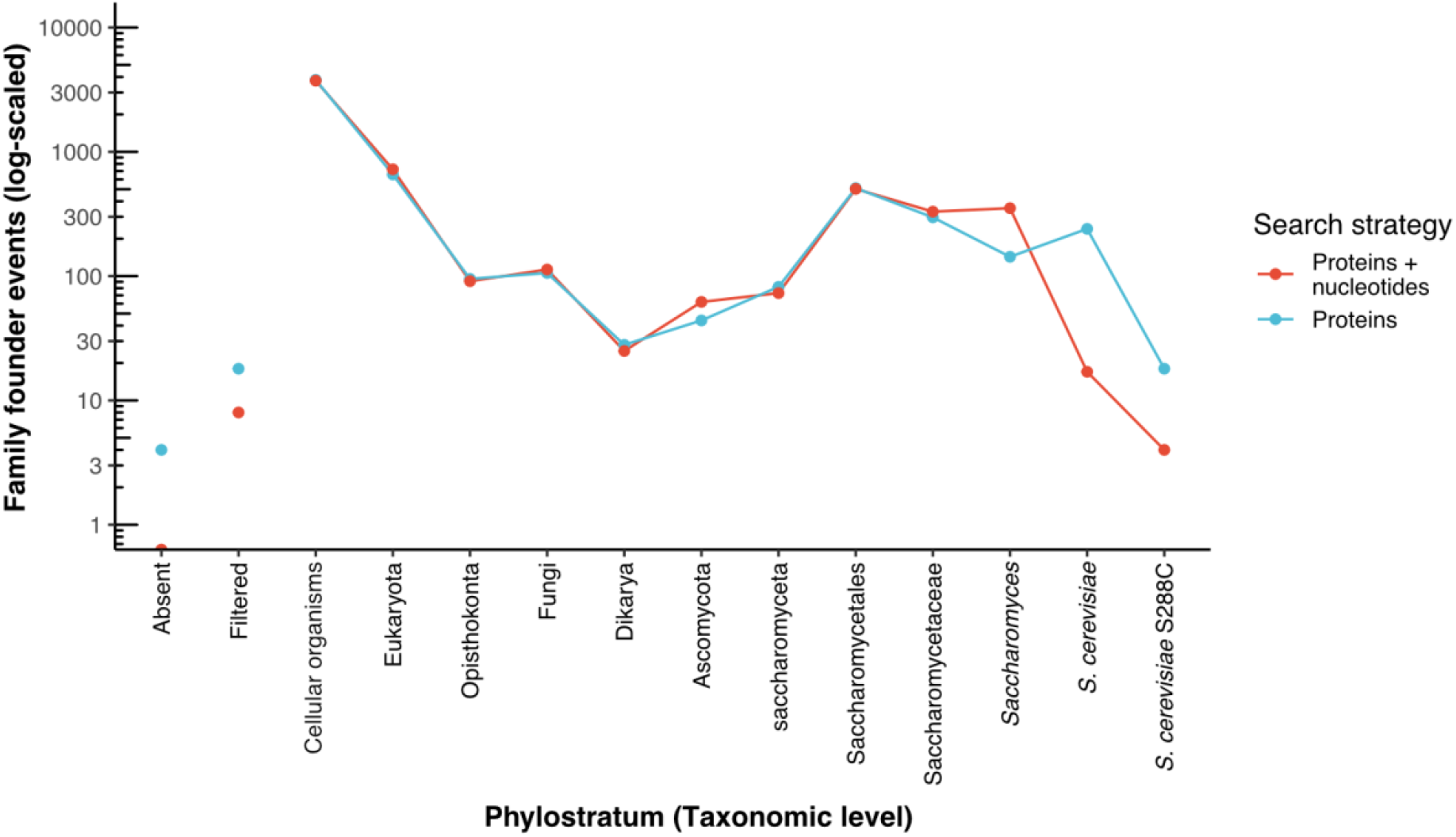
Impact of using a protein-vs-nucleotide search in addition to predicted gene annotations for the gene age assignments of *S. cerevisiae*. The standard gene age assignment was performed against the predicted genes in the NR, while the protein-vs-nucleotide search includes additional six-frame alignments against 8 representative genome assemblies from each taxonomic level, adding to a total of 80 genomes (see Table S1). The age assignment of the youngest genes are pushed to older taxonomic levels, suggesting that young gene age assignments can be overestimated when not taking annotation errors into account. However, older gene age assignments remain largely unaffected by annotation errors, demonstrating that protein-vs-nucleotide searches are mostly impactful over recent gene-family founder events. The taxonomic representativeness also improves when doing a protein-vs-nucleotide search, reducing the number of filtered gene age assignments.

**Figure S4.**
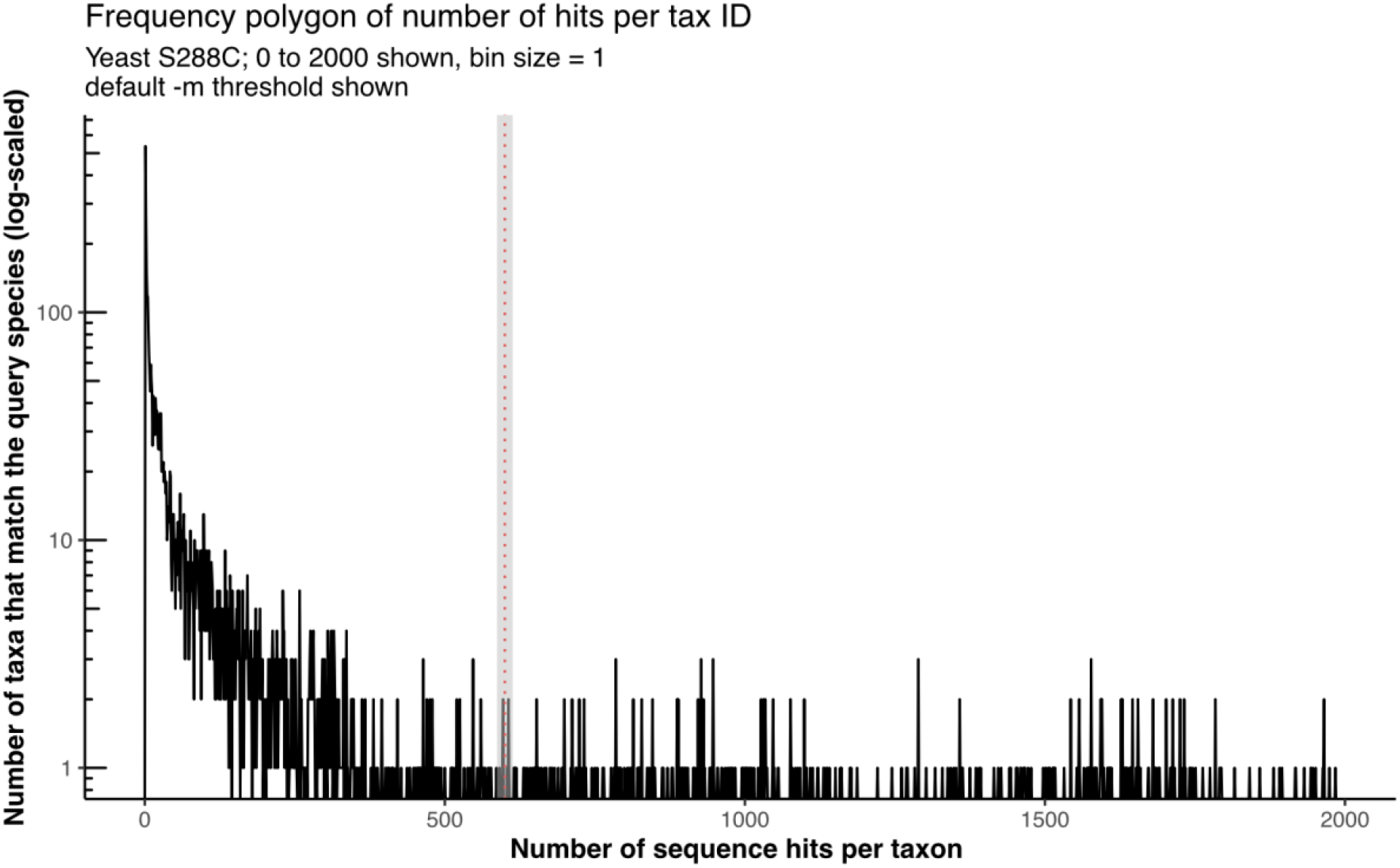
Distribution of taxa in the NR with sequence matches against the proteome of *S. cerevisiae*. Most of the taxa in the NCBI non-redundant database contain 50 or less proteins (*e*.*g*., sequence data generated for phylogenetic studies), which could lead to an unreliable gene age assignment. An empirical threshold of 10% total hits against the query proteome can successfully detect these cases (red dashed line). This allows GenEra to assign ages only to the taxonomic levels where genomic data is available.

**Figure S5.**
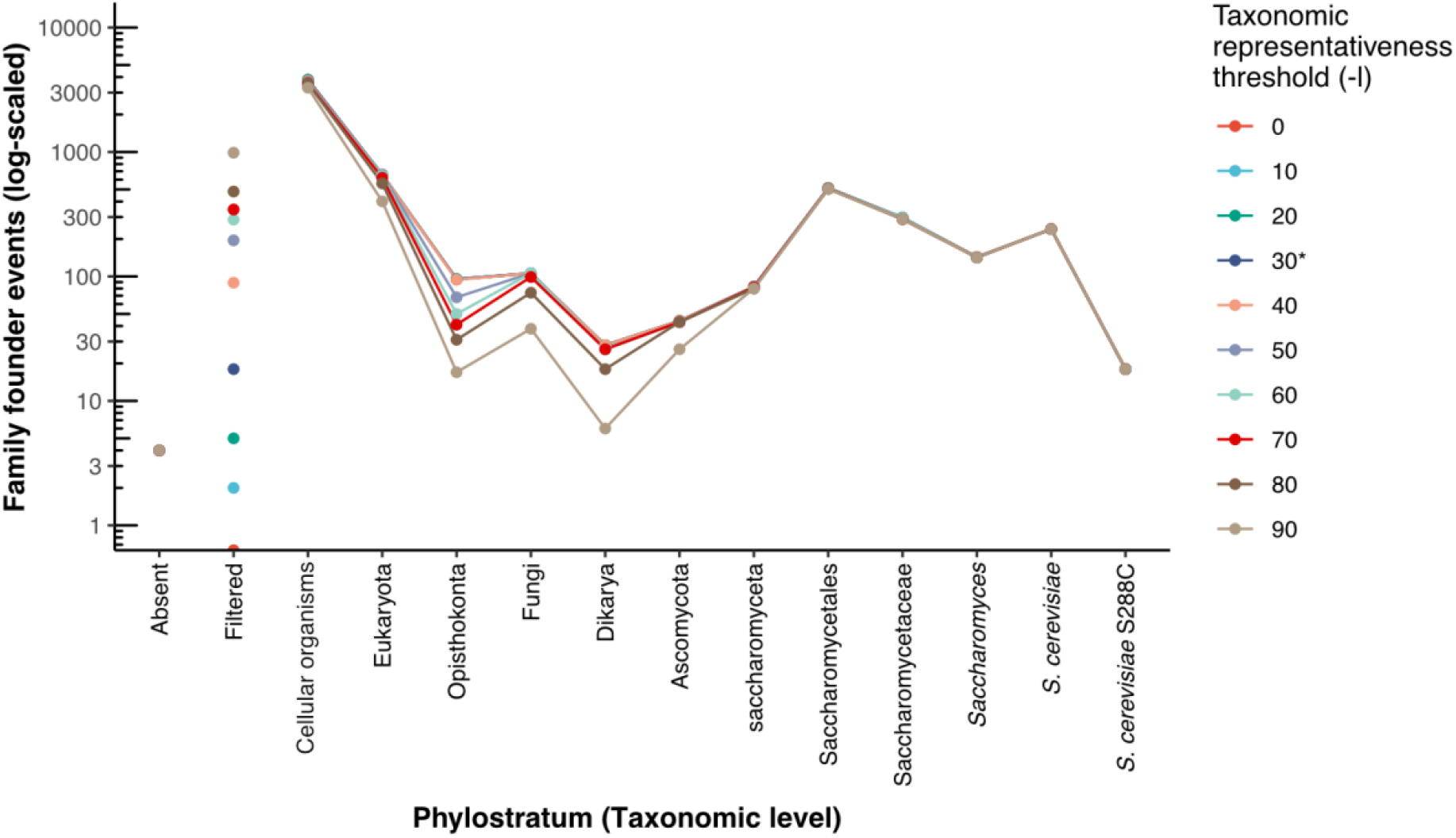
Assessment of different thresholds for the taxonomic representativeness score in *S. cerevisiae*. Higher thresholds have a direct impact on the amount of genes that cannot be assigned to a specific age (filtered). We established a default threshold of 30% for gene age assignment in GenEra, as lower values of taxonomic representativeness are bound to represent artifacts due to genome contamination or false positive matches, while establishing a more stringent threshold would fail to account for gene loss events, influencing the overall trends of gene age assignments. We established a taxonomic representativeness of 30% as the default threshold to filter ambiguous gene age assignments (asterisk).

**Figure S6.**
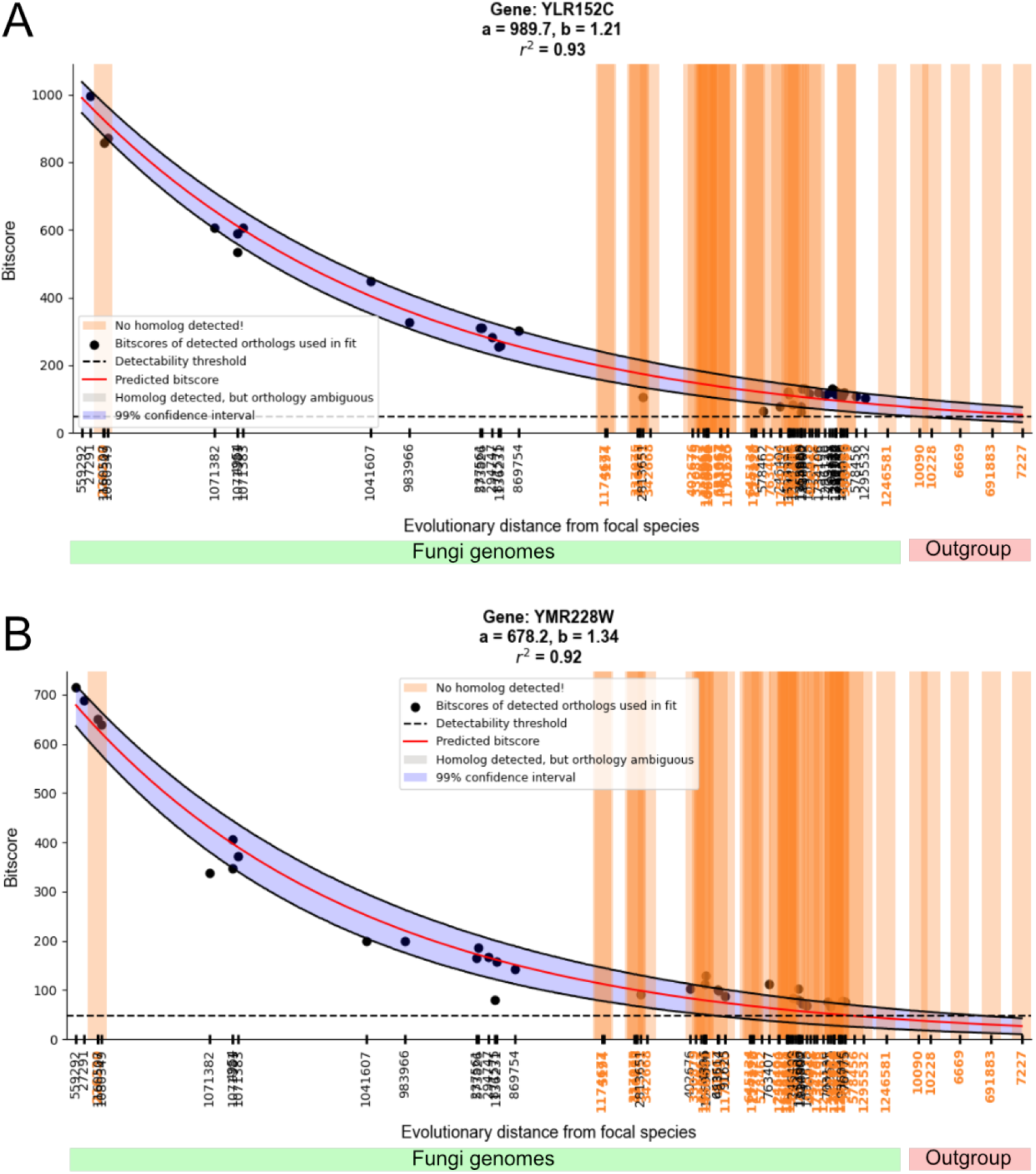
Decoupling high-confidence gene age assignments from HDF. Bitscore decay plots as a function of evolutionary distance in two genes of *S. cerevisiae* across different fungal species (green bar) and other closely related opisthokonts (red bar). Each species in the horizontal axis is represented by its NCBI Taxonomy ID. The bitscore values were retrieved from a DIAMOND search against the NR using GenEra, while the pairwise evolutionary distances (substitutions per site) were retrieved from a previously published maximum likelihood tree (43). The bitscore prediction and the detection failure probabilities were calculated using abSENSE (15). A) Bitscore decay of the gene YLR152C, which shows homolog genes across the Fungi kingdom and is predicted to have detectable bitscore values in *Fonticula alba* and in other closely-related animal genomes (detection failure probability of 0.01 in the outgroup species). This gene is thereby regarded as a high-confidence fungal TRG. B) Bitscore decay of the gene YMR228W, which is found within the Fungi kingdom, but whose expected bitscore falls below the detectability threshold (dashed line) in *Fonticula alba* and in other closely-related animal genomes (detection failure probability of 1 in the outgroup species). Therefore, the seeming absence of this gene outside the Fungi kingdom can be explained by HDF.

